# Allelic bias contributes to heterogeneous phenotypes of NK cell deficiency

**DOI:** 10.1101/2023.09.25.559149

**Authors:** Seungmae Seo, Yong-Oon Ahn, Sagar L. Patil, Jacqueline Armetta, Madrikha D. Saturne, Everardo Hegewisch-Solloa, Nicole C. Guilz, Achchhe Patel, Barbara Corneo, Malgorzata Borowiak, Emily M. Mace

## Abstract

While monogenic variants in CDC45-MCM-GINS (CMG) replisome proteins cause human natural killer cell deficiencies (NKD), family members with the same inherited variants often have variable clinical and cellular phenotypes. We investigated two siblings with inherited compound heterozygous *GINS4* variants but variable disease expressivity. Cell cycle impairment and increased apoptosis were detected following NK cell lineage commitment but not in pluripotent cells. While this effect was detected in both siblings, the efficiency of NK cell differentiation was variable and correlated with differential clinical severity of NKD. Further investigation of allelic expression of inherited GINS4 variants demonstrated expected biallelic expression of *GINS4* in pluripotent cells and progenitors. However, allelic bias in lineage-committed NK cells led to over- or under-representation of more damaging GINS4 heterozygous variants associated with differential cellular and clinical severity. This study identifies allelic bias that causes phenotypic variation of monogenic diseases and defines mechanisms underlying immunodeficiency.

## INTRODUCTION

More than 450 inborn errors of immunity have been described as arising from inherited germline genetic variants^1^. Despite demonstrated monogenic origins, many of these lead to variable expressivity of clinical and cellular phenotypes that has previously been ascribed to modifier genes, environmental exposures, or somatic reversion^2–6^.

Mutations in components of the CDC45-MCM-GINS (CMG) helicase complex, including MCM4, MCM10, GINS1, and GINS4, are associated with human diseases including immunodeficiency, growth retardation, adrenal insufficiency, Meier-Gorlin syndrome, and cardiomyopathy^7–16^. As the CMG complex is ubiquitously expressed and required for eukaryotic DNA replication, deletion of any of its core subunits causes embryonic lethality^17–20^. Inherited partial loss-of-function variants often lead to immunodeficiency that includes decreased frequencies of mature NK cells and accompanying NK cell dysfunction^7–11^. In some cases, neutropenia or T cell dysregulation has been reported, however NK cell deficiency (NKD) has been described as the predominant immune deficiency in individuals with *GINS1, GINS4, MCM10, CDC45,* and *MCM4* variants. While there is some association between the degree of damage of individual CMG variants and clinical severity, there is also a heterogeneity of disease even between siblings carrying the same variants, as noted previously for disease-causing GINS4 and MCM4 variants^7,10^. The unique sensitivity of NK cells and, in some cases, neutrophils to these variants is still incompletely understood, but likely arises from cell lineage-specific differences in sensitivity to mild replication stress that is generated by partial loss-of-function variants^21,22^. As such, understanding how immune cell lineages regulate the expression and function of replisome proteins and responses to decreased replisome function will provide important understanding into the origin of disease in these individuals and the cell biology of immune cells.

Here, we sought to better understand the variability of NKD by studying siblings with compound heterozygous variants in *GINS4* with variable disease expressivity. Using iPSCs, we identified a proliferative stage of NK cell development following lineage commitment that is specifically vulnerable to loss of GINS4 function. We further demonstrated that allelic bias results in reproducible differential expression of *GINS4* variants in NK cells but not stem and progenitor cells. This allelic bias led to a greater impairment of NK cell development *in vitro* and *in vivo* despite cells sharing the same genotype. Together, our study defines how partial loss-of-function GINS4 variants cause selective NK cell deficiency and identifies the potential influence of allelic bias on human immunodeficiency.

## RESULTS

### Variable cellular and clinical phenotypes in siblings with shared GINS4 variants

Apparently healthy parents are heterozygous for single *GINS4* variants (Fig. 1A). The mother’s (I.1) c.571C>T variant creates a premature stop codon at p.191 (p.Q191X) that leads to protein instability, and the father’s (I.2) c.511G>C variant is a missense mutation (c.511G>C, p.V171L) ^10^ (Fig 1B). Their children (II.1 and II.2) have compound heterozygous variants resulting in significantly decreased levels of GINS4 protein^10^. Despite the same genotype, II.1 has more severe and frequent viral infections than II.2^10^ (Fig. 1C). Repeated flow cytometric analyses of peripheral blood demonstrated that this clinical phenotype is associated with a circulating NK cell phenotype characterized by decreased NK cell number and lower frequency of mature CD62L^neg^ CD16^+^ CD56^dim^ NK cells in II.1 compared to II.2^10^ (Fig 1D). Similarly, despite not having clinical manifestations of immunodeficiency, the WT/Q191X heterozygous carrier (I.1) had decreased frequencies of mature NK cells relative to healthy controls and II.2 associated with greater instability of GINS4 protein^10^.

**Figure 1.**
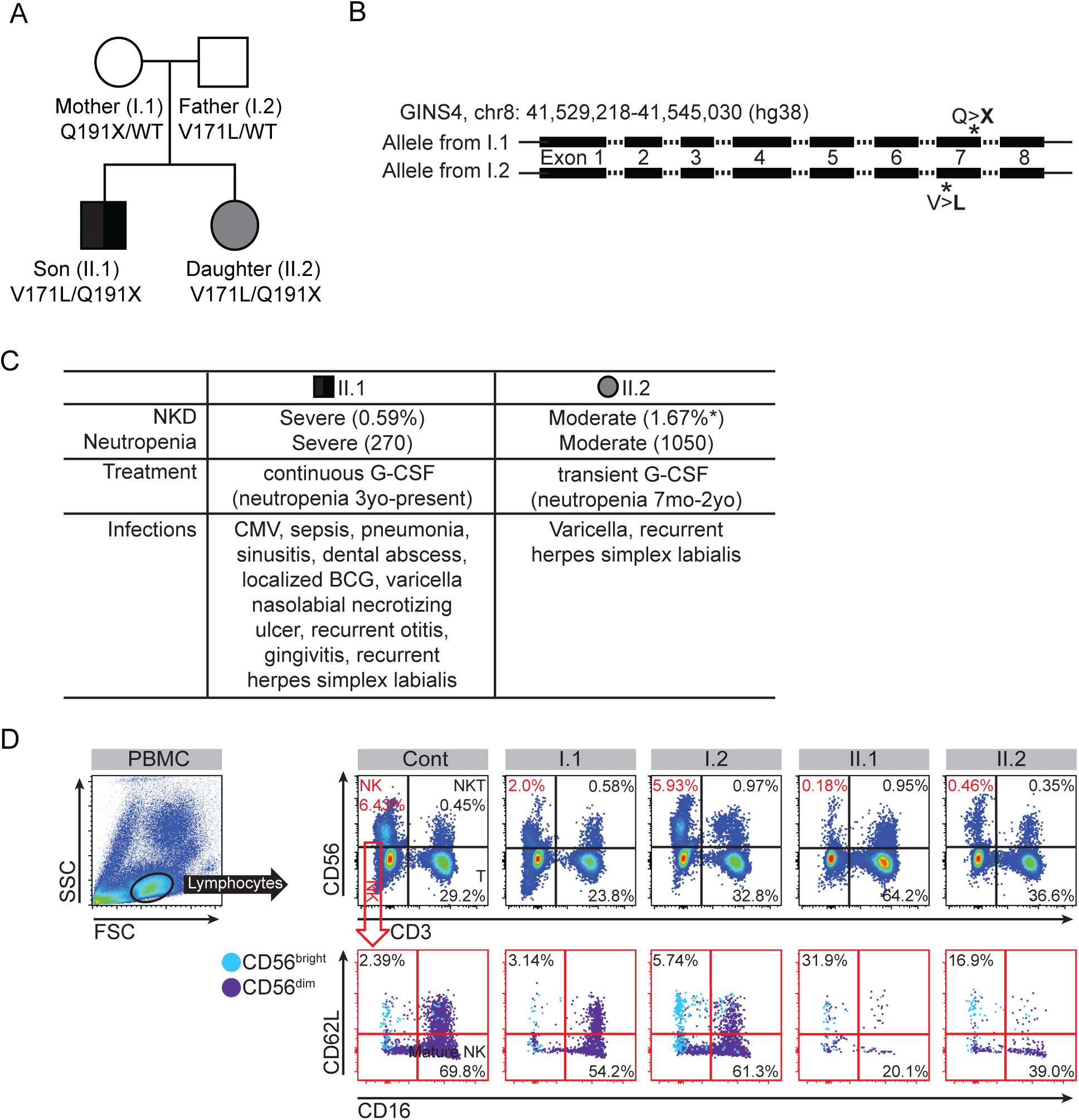
Compound heterozygous GINS4 variants result in NKD with incomplete penetrance. **(A)** Pedigree showing *GINS4* variants in the family described in ^10^. **(B)** Structure of *GINS4* gene and location of the variants. **(C)** Summary of the clinical phenotype of II.1 and II.2 individuals (n=4, student t-test *p<0.05, compared to II.1 NK cell frequency). **(D)** Dot plots showing representative flow cytometry data of peripheral blood demonstrating NK cell frequency (top) and phenotype (bottom). Frequencies of mature NK cells (CD56+CD16+CD62L-) and immature NK cells (CD56+CD16-CD62L+) are shown. n=4.

### Generation of immune and non-immune cells from pluripotent stem cells

iPSC lines were generated from PBMCs from parents (I.1, I.2), siblings (II.1, II.2) and an unrelated healthy control (C.1). Following reprogramming and validation of pluripotency, we corrected both alleles of *GINS4* in the II.1 line to generate an isogenic control line (II.1-Cor). Differentiation was performed using the spin-EB method, which supports NK cell differentiation by generating autologous stromal cells together with hematopoietic progenitors which subsequently give rise to mature, functional NK cells^23^. On day 6 of differentiation, CD34^+^ endothelial and hematopoietic precursors emerged and CD34^−^ stromal cell precursors were present (Fig. 2A). Subsequently, CD34 expression decreased as hematopoietic cells sequentially gained CD45, then CD56, CD117, CD94, and granzyme B, indicating NK cell specification that was robust by day 28 and functional maturation evident by days 35-42 of differentiation (Fig. 2A, Panel 1, Table S1).

**Figure 2.**
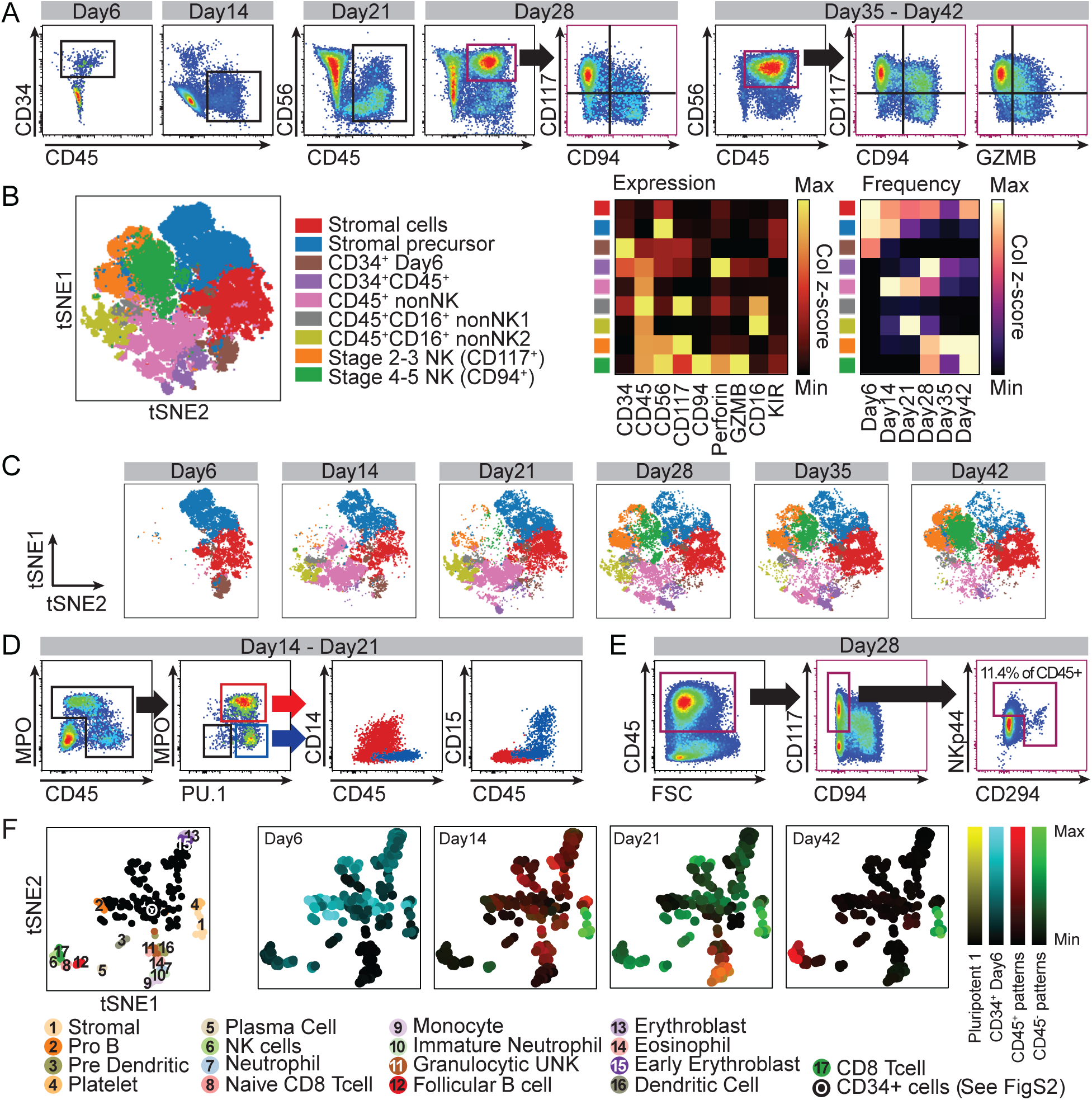
Trajectories of NK cell emergence from iPSCs. Six iPSC lines (*GINS4* variant carriers: I.1, I.2, *GINS4* variant lines: II.1, II.2, an unrelated healthy control (C.1), and an isogenic control (II.1-Cor)) were differentiated to NK cells with autologous stroma. Flow cytometry was performed to monitor the progression of NK cell differentiation, and the data were concatenated, normalized, and visualized. **(A)** Dot plots showing a representative flow cytometry result of the control (C.1) line. CD34^+^ cells emerge on day 6. Hematopoietic lineage cells start expressing CD45 on day 14, and CD56, CD117, CD94, and granzyme B (GZMB) emerge sequentially between days 21 and 42. Not shown: FSC/SSC and doublet exclusion. **(B)** tSNE plot summarizing the flow cytometry data of a representative experiment from all six cell lines at 6 weekly time points (n=5-13 technical replicates). Heatmaps show a summary of relative protein expression in the nine clusters (left) and temporal changes of each cluster’s frequency (% of cells in each cluster from all live cells; right). **(C)** Parsed tSNE showing the changes in cell populations from days 6 to 42. **(D)** Representative flow cytometry results showing the myeloid identity of non-NK cell CD45^+^ populations on days 14 and 21 (n=3 technical replicates). The gating strategy shows selection of CD45^dim^ and CD45^+^ populations. MPO^+^CD45^dim^PU.1^+^ cells in red box (middle) are represented as red dots in CD14 by CD15 plots (right), and MPO^−^CD45^+^PU.1^+^ cells in blue box (middle) are represented as blue dots in CD14 by CD15 plots (right). **(E)** Representative flow cytometry results showing non-NK ILCs in CD45^+^ populations on day 28. Gating strategy shows the selection of CD45^+^CD117^+^CD94^−^ population. The frequency of NKp44^+^ or CD294^+^ ILC-like cells are calculated as % of CD45^+^ cells. **(F)** Projection of the timepoint- and condition-specific CoGAPS patterns onto human bone marrow atlas data ^29^. Left: tSNE with the original cell type annotation. Right: Projection of CoGAPS patterns. Color intensities represent higher enrichment of each pattern.

We first sought to generate the landscape of NK cell emergence in concert with the other cell types present in spin-EB differentiation cultures. Flow cytometry data were concatenated from all donors and weekly timepoints then normalized and visualized following tSNE dimensionality reduction (Fig. 2B). Using normalized expression of these markers we identified nine populations, including two CD34^−^CD45^−^ stromal cell populations, one CD34^+^ cluster present at day 6, one CD34^+^CD45^+^ cluster, two CD45^+^CD16^+^ non-NK clusters, and two NK cell clusters. The NK cell clusters corresponded to immature NK cell developmental stages that expressed CD56 and CD117 (stages 2-3), and mature NK cells with CD56, CD94, perforin, and granzyme B (stages 4-5)^24^ (Fig. 2B, Fig. S1A). NK cell maturation marked by CD94, CD117, perforin, and granzyme B expression was prominent by day 35 and maintained through day 42 (Fig. 2B, C).

We also wanted to further characterize the CD45^+^CD16^+^ non-NK cell clusters that were transiently present at days 14 and 21. Additional flow cytometry analysis showed that these were MPO^+^CD45^dim^ PU.1^+^CD16b^+^CD15^+^ granulocytes and MPO^−^CD45^+^PU.1^+^M-CSFR^+^CD32^+^CD14^+^ monocytes (Fig. 2D, Fig. S1B, Table S1, Panel 2). In addition, NKp44 and CD294 staining identified 10-15% of CD45^+^ cells on days 28 and 35 to be CD117^+^CD94^−^, CD294^+^ or NKp44^+^ ILC-like cells (Fig. 2E, S1C, Table S1 Panel 3)^25–27^. Notably, while the number of CD45^+^ cells at timepoints associated with NK cell generation were reduced in II.1 and II.2 lines (discussed further below), these myeloid and ILC-like cells were present in all lines at similar frequencies.

To better understand NK differentiation using transcriptome-wide signatures, we performed bulk RNA-seq on iPSCs, sorted CD34^+^ cells on day 6, and separated CD45^+^ hematopoietic and CD45^−^ non-hematopoietic/mesodermal cells on days 14, 21, and 42 of differentiation. A Bayesian non-negative matrix factorization method using Coordinated Gene Activity in Pattern Sets (CoGAPS) defined 15 transcriptional modules that shape NK cell emergence^28^. Of the 15 CoGAPS patterns, nine patterns were time- and cell type-specific (Fig. S1D, S1E, Table S2, S3). A pattern that represents day 42 CD45^+^ samples was enriched for NK cell markers (Table S2). We noted that expression levels of the top genes in this pattern were significantly lower in the CD45+ cells isolated from II.1 at day 42. Interestingly, we found that this NK cell specific gene signature was also decreased in CD45^+^ cells from the I.1 line, whereas this signature was present in the II.2 line at similar levels to I.2, C.1, and the corrected line (II.1-Cor) (Fig S1F).

Finally, to align the transcriptomic data generated from in vitro differentiation to in vivo data, we projected these CoGAPS patterns to the summarized human bone marrow cell atlas scRNA-Seq data cataloging 35 cell types from 8 individuals^29^. The CD34^+^ day 6 pattern aligned most closely with CD34^+^ progenitor populations, specifically HSCs and lymphoid-biased cells (pre-B cycling, CLP, HSC, HSC cycling, pre-B, and unknown lymphoid) (Fig. 2F, S1G). The CD45^+^ day 14 pattern aligned with CD34^+^ megakaryocyte progenitor and CD34^+^ granulocyte progenitors. As predicted by flow cytometry data, the CD45^+^ day 21 signature projected to granulocytes, with the most enrichment in the neutrophil, immature neutrophil, and monocyte subsets (Fig. 2D, F). Finally, the CD45^+^ day 42 signature was enriched in NK cells (Fig. 2F). CD45^−^ signatures from our data consistently aligned with the stromal cell cluster present in the bone marrow dataset, further confirming their non-hemogenic identities (Fig 2F).

### Variable efficiency of NK cell generation from GINS4 variant iPSCs

As suggested by the RNA-seq data, tSNE projections of flow cytometry data from repeated differentiation experiments revealed a lower efficiency of NK cell generation from the II.1 than the II.2 line (Fig 3A, B, S2A). In contrast, the II.1-Cor and C.1 control lines reproducibly generated mature NK cells. To better understand experimental and cell-line-dependent variation^30,31^, we included two more unrelated control iPSC lines (C.2 and C.3) in our differentiation experiments. In >50% of our technical replicates in the three unrelated controls (C.1, C.2, C.3), the parent lines (I.1 and I.2), and the II.1-Cor line generated CD45^+^CD56^+^ NK cells at >30% frequency by day 35 (Fig 3B, S2B, n=5-13). 67% of differentiation experiments from II.2 cells resulted in >30% CD45^+^CD56^+^ NK cells, which was within the range of normal control line variation. In contrast, only 23% of experiments differentiating the II.1 line resulted in >30% NK cells (Fig 3B). Further, in cases when NK cells were generated, II.2 CD45^+^CD56^+^ NK cells had a more mature phenotype, with a higher frequency of CD94^+^ and granzyme B^+^ cells than those generated from II.1 (Fig 3C). Importantly, no differences in lineage potential were detected by colony-forming assays or T cell differentiation to CD4/CD8 double-positive cells, confirming the NK cell-specific disease phenotype (Fig 3D, S2C).

**Figure 3.**
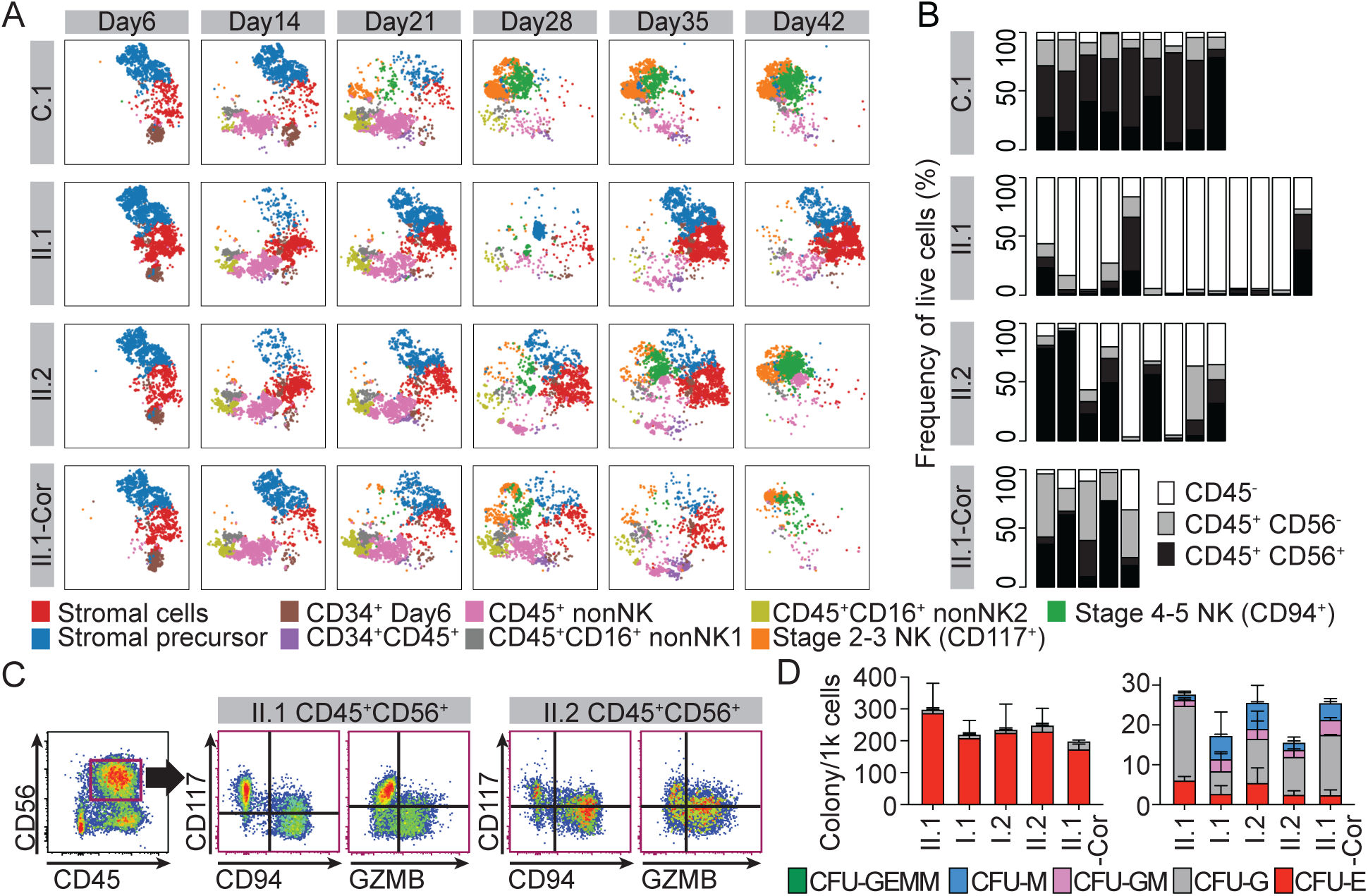
iPSC lines from II.1 and II.2 have variable NK potential. All differentiation experiments were summarized to show the cell line specific efficiency for NK cell differentiation in each line. **(A)** Parsed tSNE plots showing changes in cell populations in lines C.1, II.1, II.2, and II.1-Cor from days 6 to 42. **(B)** Stacked bar graphs showing the population frequencies of cells with hematopoietic (CD45) and NK cell (CD56) markers on days 35 or 42 of each experiment from C.1, II.1, II.2, and II.1-Cor lines (n=5-13). Each bar represents a single experimental repeat, and the frequencies are calculated as % of live single cells. **(C)** Representative flow cytometry dot plot showing the expression of NK cell markers, CD94, CD117, and granzyme B in CD45^+^CD56^+^ cells from experiments where the II.1 and II.2 lines generated NK cells at the highest frequency. **(D)** Colony forming assays showing the lineage potential of each cell line when modeling primitive hematopoiesis (left) or definitive hematopoiesis (right) (n=3-9 technical replicates, error bars = SD).

Given the presence of autologous stroma in our differentiation conditions, we sought to rule out the contribution of differences between stromal cells to our phenotype. We sorted iPSC-derived day 6 CD34^+^ cells and differentiated these on OP9 stromal cells ^32^. In all lines tested, we found that this protocol was less efficient at generating NK cells, likely due to the separation of progenitors from stromal cells. Under these less efficient culture conditions, both II.1 and II.2 lines were equally inefficient in generating NK cells relative to C.1, C.2, C.3, I.1, I.2, and II.1-Cor lines (Fig S2D-I)^23^. Together, these data demonstrate the capacity of II.2 for improved NK cell generation under optimized conditions, whereas II.1 was consistently impaired despite sharing the same compound heterozygous *GINS4* genotype.

### GINS4 deficiency disrupts cell cycle progression in committed NK cells

We next wanted to investigate the functional impact of GINS4 variants on cell cycle progression and cell viability. There was significantly lower GINS4 protein expression in 11.1 and II.2 pluripotent cells than control and parent lines, consistent with our previous studies^10^, but GINS4 protein levels were rescued in the isogenic control line (II.1-Cor). This pattern was consistent up to day 14, when the difference became insignificant due to decreased expression of GINS4 in all cell lines as cells underwent differentiation to the NK cell lineage (Fig. 4A, Fig. S3A). Expression of other CMG helicase complex components, including MCM4 and MCM6, were unaffected in the II.1 and II.2 lines (Fig. S2A, S3B). Unlike protein, GINS4 mRNA levels in the II.1 and II.2 lines were equivalent to the control lines, supporting the instability of the variant protein, but not the RNA (Fig. 4B)^10^.

**Figure 4.**
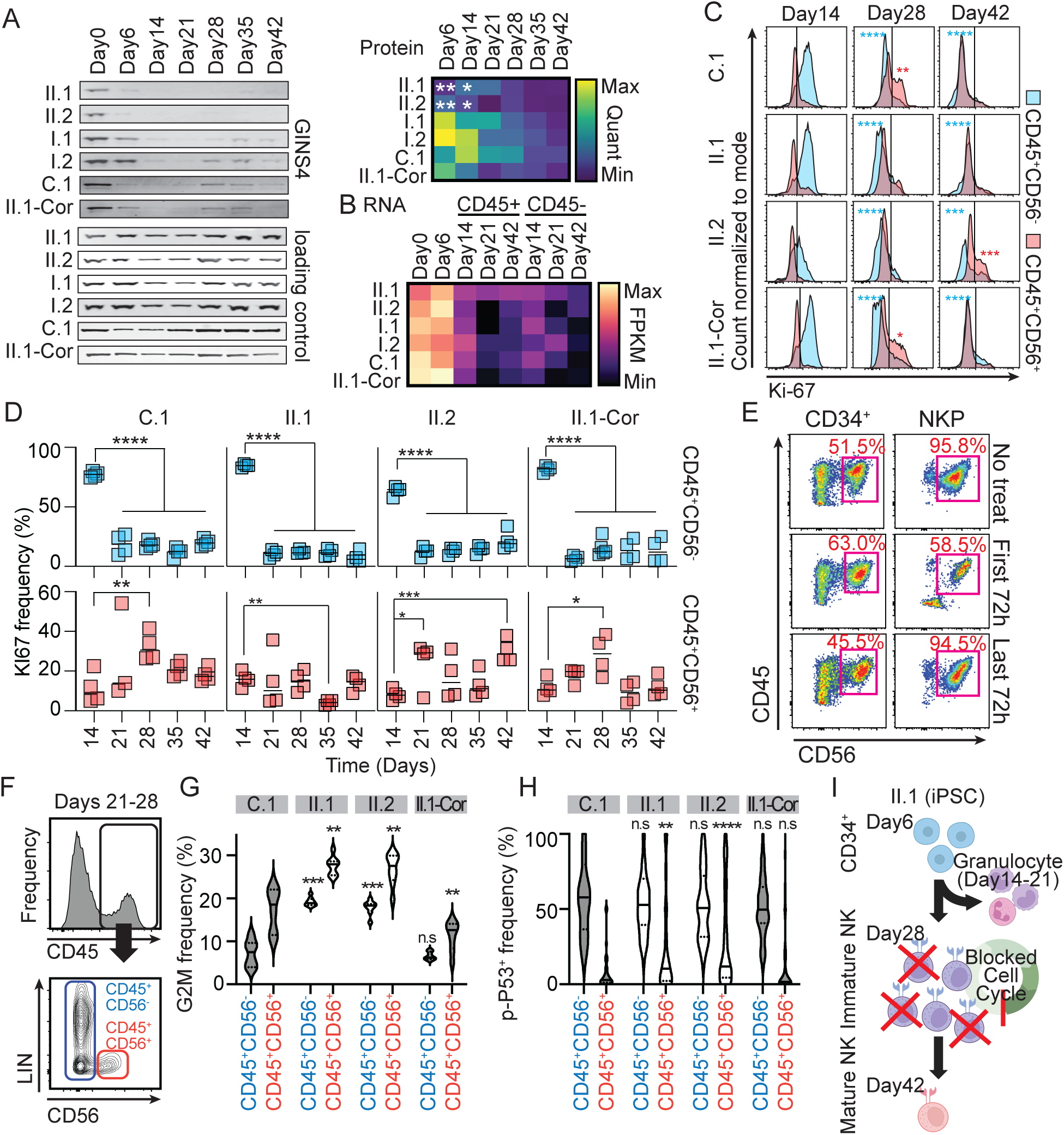
GINS4 deficiency disrupts the cell cycle progression of committed NK cell progenitors. Cell cycle and apoptosis signals were measured during NK differentiation from GINS4 variant iPSC lines. **(A)** Western blot for GINS4 in iPSCs (day 0) and throughout NK cell differentiation. Heatmap shows averaged normalized expression of GINS4 from Western blots (n=3, student t-test *p<0.05, **p<0.01). **(B)** Heatmap shows transcriptional expression of *GINS4* from RNAseq. **(C)** Representative flow cytometry histograms showing Ki-67 levels of CD45^+^CD56^-^ (blue) and CD45^+^CD56^+^ (red) populations in C.1, II.1, II.2, and II.1-Cor lines at days 14, 28, and 42 of differentiation. **(D)** Quantification of the frequency of CD45^+^CD56^−^Ki-67^+^ (top), and CD45^+^CD56^+^Ki-67^+^ (bottom) populations in C.1, II.1, II.2, and II.1-Cor lines at all time points (n=4 technical replicates, student t-test *<0.05, **<0.01, horizontal lines indicate average). **(E)** Representative flow cytometry results showing the differentiation efficiency of immature NK cells (CD45^+^CD3^-^CD117^+^ lymphocytes) from cord blood under replication stress. Cells were treated with 0.25 µM aphidicolin for the first 72 hours or the last 72 hours of 14 days of differentiation (n=3). Mature NK cells are defined as CD45^+^CD56^+^ cells, and the frequencies are calculated as % of live single cells. **(F)** Representative flow cytometry data showing CD45, lineage (Lin; CD13, CD14, and CD15), and CD56 population gating. **(G)** Violin plot showing cell cycle profiles quantified from DAPI in each population represented in (F) from C.1, II.1, II.2, and II.1-Cor lines (n=8 technical replicates, student t-test **p<0.1, ***p<0.005, p-values represent G2/M frequencies compared to the same population in C.1 line). **(H)** Frequency of phospho-p53 (Ser15)^+^ cells in each population from days 21 to 28 combined from C.1, II.1, II.2, and II.1-Cor lines (n=6-9 technical replicates, student t-test **p<0.01, ****p<0.001, p-values represent p-P53^+^ frequencies compared to the same population in C.1 line). **(I)** Model showing a proposed mechanism of pathogenesis of NKD (in red) resulting from impaired GINS4 expression.

Despite the significant difference in GINS4 protein expression, no differences in cell cycle progression between lines were observed in pluripotent cells or at day 6 (Fig. S3C, S3D). Hematopoietic non-NK cells (myeloid cells) at day 14 were robustly proliferative from all cell lines. Following NK cell commitment, we detected subpopulations of CD45^+^CD56^+^ NK progenitor cells in the control lines that were proliferating, particularly at day 28 (Fig. 4C, D). CD45^+^CD56^+^ cells generated from II.1 line consistently failed to robustly proliferate; like our immunophenotyping data, while 11.1 line was more variable and had a reproducibly delayed proliferative phase seen at day 42 (Fig. 4C, D).

These data suggested that proliferation following NK cell lineage commitment and overall decreased expression of replisome proteins contribute to a unique sensitivity of NK cells to replication stress. To test how mild replication stress affects different stages of NK cell development, we isolated CD34^+^ hematopoietic progenitors and CD117^+^ NK cell progenitors^17^ from healthy donor cord blood and differentiated them into mature NK cells in the presence or absence of transient aphidicolin-induced replication stress to mimic the effects of loss of CMG helicase function. NK progenitors in the first 72 hours of differentiation were more sensitive to aphidicolin treatment than CD34^+^ cells or cells that had completed maturation, underscoring lineage commitment to conventional NK cells as being vulnerable to replication stress (Fig 4E).

To better understand the effect of GINS4 variants on cell cycle progression, we measured DNA replication every 24 hours during the emergence of CD45^+^CD56^+^ NK cells between days 21 and 28 (Fig. 4F-H). Compared to the II.1-Cor line, II.1 and II.2 CD45^+^ cells, especially CD45^+^CD56^+^ NK cells, were more frequently found in G2/M phase, particularly after commitment to the NK cell lineage (Fig. 4G, Table S1 Panel 4). This finding is consistent with the relatively mild cell cycle phenotype leading to G2/M accumulation caused by these GINS4 variants^10^. We also recently demonstrated that primary NK cells are more sensitive to replication stress relative to T cells^22^. CD45^+^CD56^+^ NK precursor cells from II.1 and II.2 also had significantly higher frequencies of phospho-p53 (Ser15)^+^ cells than II.1-Cor line (Fig. 4H, Table S1 Panel 5). These results suggest that replication stress shortly after NK cell lineage commitment specifically impairs NK cell due to increased apoptotic activity (Fig. 4I).

### Allelic bias contributes to variable expressivity of *GINS4* variants

We repeatedly found that despite the same GINS4 genotype, II.1 and II.2-derived cells had differences in their functional and phenotypic responses. Correction of both alleles in the II.1 line to reference alleles corrected all tested phenotypes of the II.1 lines, demonstrating that the *GINS4* gene was responsible for these effects. A recent study on inborn errors of immunity (IEI) has implicated seemingly random monoallelic gene expression as a cause of incomplete penetrance of IEI genes^33^. Given reproducible differences in clinical phenotypes that were consistent with the incomplete penetrance of NKD, we tested whether allelic bias may account for differential usage of the Q191X or less damaging V171L allele. We isolated primary NK (CD56^+^CD3^-^), T (CD3^+^), and B (CD19^+^) cells from the parents (I.1, I.2) and siblings (II.1, II.2) and performed bulk RNA-seq (Fig 5A). Due to the reduced numbers of NK cells in the peripheral blood of II.1 and II.2, we pooled PBMC samples from 2-3 collection dates. Consistent with previous flow cytometry data (Fig. 1), primary NK cells from II.1 had lower expression of NK cell maturation markers relative to the control and parents, while II.2 cells showed equivalent levels of expression (Fig. 5B). Given the non-proliferative state of resting blood cells and low *GINS4* expression in differentiated cells, *GINS4* expression was too low in the NK cells to assess ‘transcriptotypes’ by RNA-Seq. Therefore, we performed digital PCR to amplify *GINS4* transcripts in these samples. A silent mutation in *NOC2L* was used as a technical control, and allelic bias of *NOC2L* expression was not observed in any of our samples (48-51%, Fig. S3E).

**Figure 5.**
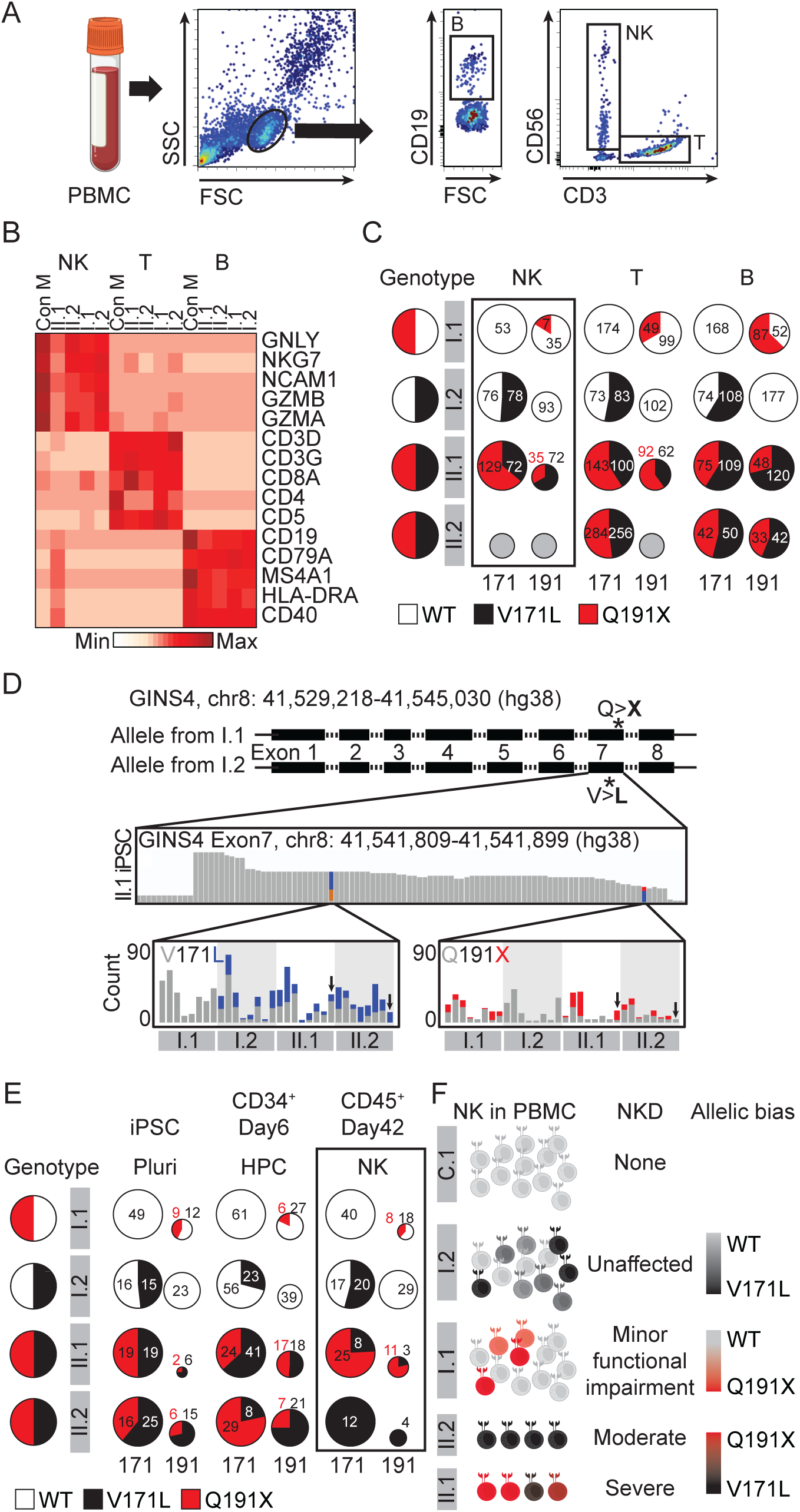
Allelic bias of GINS4 expression is in II.1 and II.2 lines. Sorted PBMC lymphocytes and iPSC-derived NK cells were assessed for transcriptional allelic bias. **(A)** Representative dot plots show the sorting strategy for B, NK, and T cells from PBMC. Lymphocytes were gated on FSC/SSC and CD19, CD56, and CD3 were used to identify cell populations. **(B)** Heatmap of NK, T, and B cell markers confirms subtype identity in control (Con M), and GINS4 family member (I.1, I.2, II.1, II.2) PBMC samples. **(C)** Digital PCR result represented in pie charts show the contribution of each allele within sorted lymphocytes. Size of the pie charts represent relative concentration of the detected p.191 compared to p.171 locus in each sample. Numbers in the pie charts show concentration (copy/μl) per allele. Gray circles represent no detection. **(D)** Structure of the *GINS4* gene (top) and IGV track showing the frequencies of V171L and Q191X variants from II.1 iPSC RNA-seq sample (middle). Bar graphs (bottom) showing the frequency and ratio of V171L and Q191X variants in all iPSC derived RNA-seq samples. Arrows indicate day42 CD45^+^ samples of II.1 and II.2 lines. **(E)** Pie charts showing the contribution of each allele in all family member lines at pluripotency, day 6 CD34^+^, and day 42 CD45^+^ cells. Size of the pie charts represent relative counts at the p,191 locus compared to p.171 locus in each sample. Numbers in the pie charts show counts per allele. **(F)** A model showing the connection between the GINS4 ‘transcriptotype’ and NKD severity in all family members. Color range shows allelic bias in gene expression.

Unfortunately, we were unable to amplify *GINS4* from the RNA sample from II.2 NK cells. NK cells from II.1 showed distinct bias towards expression of the mother’s and father’s allele at p.171 locus and p.191 locus, respectively (Fig. 5C). Considering the low expression and instability of the variant, together with the C-terminal proximity of p.191, we suspect the p.171 locus is a better measure of allelic bias. However, since replication with primary cells was challenging due to limited availability of primary samples, we sought to determine if allelic bias could be measured in iPSC-derived cultures. In our bulk RNA-seq data from all donors and timepoints throughout NK cell differentiation (Fig. 2), we observed unequal expression of *GINS4* alleles in multiple samples, suggesting *GINS4* is frequently subject to allelic bias (Fig. 5D-E, S3F). The p.191 locus consistently had poorer coverage compared to the p.171 locus (Fig 5E, size of the pie charts). As seen in primary NK cells, day 42 CD45^+^ cells from I.1 were 62% biased towards the reference allele when considering the p.191 locus and I.2 was unbiased based on p.171 expression. II.1 was 77% biased towards p.Q191X allele when considering both variant loci (Fig 5E). The day 42 CD45^+^ cells from II.2 expressed the p.V171L allele 100% of the time, demonstrating a connection between variable expressivity of NKD and allelic bias in *GINS4* expression (Fig 5E). Together, these data suggest that the clinical and cellular phenotypic variability we observe between II.1 and II.2 may be due to distinct epigenetic regulation of GINS4 alleles in differentiated cells (Fig. 5F)

## DISCUSSION

CMG helicase variants have previously been described to cause NK cell deficiency with or without accompanying neutropenia or mild T cell lymphopenia, but why certain immune subsets are selectively vulnerable to these variants remained unclear^21^. Our use of patient-derived iPSC lines in this study enabled the dissection of the cell cycle dynamics of human NK cell differentiation and identified a critical proliferative stage during which CD45+CD56+ progenitors emerge. In both siblings previously shown to have NKD^10^, compound heterozygous variants in *GINS4* result in G2/M cell cycle arrest and increased apoptotic activity at this proliferative stage, identifying the reactivation of cell cycle following commitment to the NK cell lineage as being vulnerable to cell cycle impairment resulting from partial loss of GINS4 expression. As the variants in this family lead to decreased GINS4 protein expression, our study connects the decreased expression of GINS4 caused by protein instability with the intrinsic down-regulation of replisome proteins through NK cell differentiation. These deficits converge specifically on proliferative NK precursors while subsets with higher replisome protein expression, including pluripotent cells, are spared.

The use of patient-derived and isogenic control iPSCs in our study was a powerful tool for modeling a rare genetic defect from a single family. As previously demonstrated^30,31,38,39^, we observed heterogeneity between and within iPSC lines from healthy donors. However, through multiple rigorous experiments, we also recapitulated previously noted heterogeneity in cellular and disease phenotypes between siblings sharing the same GINS4 genotype^10^. Consistent with primary NK cell transcriptional and cell surface phenotypes from these individuals, iPSCs from the more severely affected individual (II.1) demonstrated inefficient NK cell generation and the NK cells that were generated had a less mature phenotype. In contrast, iPSCs from II.2, the sibling with a milder clinical presentation, exhibited greater NK cell differentiation potential despite having functional cell cycle impairment.

A recent study on inborn errors of immunity implicates seemingly random monoallelic gene expression as a cause of incomplete penetrance and variable expressivity of IEI genes^33^. Consistent with this, our experiments revealed allele-specific *GINS4* variant expression in NK cells, but not in pluripotent cells (Fig 5F). In both primary NK cells and iPSC-derived NK cells, II.1 NK cells expressed more of the Q191X allele, whereas II.2 NK cells were biased towards the less damaging V171L allele. The heterozygous carrier of Q191X allele, I.1, exhibited normal NK cell counts and functionality, but had recurrent herpes labialis and her brother died from severe gastrointestinal sepsis at 9 months of age^10^. These data further suggest that Q191X is a more damaging variant, and that the milder phenotype of II.2 may be due to decreased Q191X expression in NK cells. We also noted allelic bias of GINS4 expression in other mature immune cell subsets, including T cells, B cells and other iPSC-derived CD45^+^ cells. These data suggest that the *GINS4* gene is subject to allelic bias in mature NK cells and other differentiated immune cell subsets. Further investigations should address how prevalent the random allelic biases of autosomal genes are in different cell types. In our study, we measured expression from bulk RNA-seq and thus couldn’t confirm that GINS4 had monoallelic expression at a single cell level, except for the few times where the bias was 100%. The color scheme of our proposed model is intentionally ambiguous to reflect this limitation (Fig 5F). As both variants are in the penultimate C-terminal exon, it is very challenging to assess allelic expression using current single-cell RNA-seq technology. The mechanism by which allelic bias is established and maintained, including through reprogramming of blood to iPSCs, remains unclear.

Taken together, this work identifies how changes in cell cycle dynamics, paired with the effects of CMG variants, affect the generation of human NK cells. While T cell deficiencies are less common than NK cell deficiencies in patients with disease-causing CMG variants, a 50% reduction in MCM7 expression in human T cells exiting quiescence leads to DNA damage and cell cycle delays, but not apoptosis^34^. In contrast, mature NK cells under comparable activation and mild replication stress express less CMG protein and undergo apoptosis at a higher rate than T cells^22^. As such, we interpret the repeated identification of NK cell defects associated with these variants not as an indication that NK cells are the only subset that could be affected by these variants, but instead as an illustration of varying cell lineage-dependent requirements for CMG protein concentrations that extend to lineage-specific responses to mild replication stress. There may also be roles for CMG proteins outside of DNA replication that are affected by disease-causing CMG variants. Known other roles include the function of the MCM complex in histone transfer^35,36^ and the role of PSF-2 (GINS2) in programmed cell death resulting from asymmetric cell division during *C. elegans* development^37^. These cases illustrate an emerging area of study on the importance of replisome proteins in cell fate and pluripotency choices occurring during lineage specification. How different immune subsets, and lymphocytes in particular, can differentially regulate their requirement for the generation, localization, and function of replisome proteins is a compelling area of interest, particularly given the disease associations highlighted by individuals affected by these variants. Our study also further identifies GINS4 transcriptional control as being subject to biased allelic expression revealed by unequal expression of two inherited missense variants. While the mechanisms underlying monoallelic expression are incompletely understood, in this case, it explains incomplete disease penetrance and accompanying differences in molecular function.

## MATERIALS AND METHODS

### Primary cells and generation of iPSC lines

Peripheral blood from healthy donors and patients was collected under the guidance of the IRB (protocol #AAAR7377) at Columbia University in accordance with Declaration of Helsinki guidelines. PBMCs were isolated by density gradient centrifugation and cryopreserved. iPSCs were generated from cryopreserved PBMC from GINS4 affected individuals, their parents, ^10^ and unrelated female adult healthy donor controls using StemRNA Gen Reprogramming Kit (Stemgent) as previously described ^8^. Cells were validated for pluripotency and periodically karyotyped and tested for mycoplasma.

### Cell culture

Human iPSCs were cultured in feeder-free condition with Stemflex (Gibco, A3349401) or mTeSR plus (StemCell Technologies, 100-0276) at 37°C with 5% CO2. Cells were dissociated to single cells with Accutase (Sigma, A6964), plated at a density of 1 × 10^6^ cells/well in a Matrigel-coated 6 well plates (BD, 354277; Fisher Scientific, 140675). For the first 24 hours after passaging, cells were cultured with 5μM ROCK inhibitor (Y27632; StemCell Technologies, 72304) and passaged every five days in culture. OP9 cells (ATCC CRL-2749) were cultured in nongelatinized culture flasks (Fisher Scientific, 10-126-28) in α-MEM (Fisher Scientific, 12561056) with 20% heat-inactivated FBS (Gemini, 900-108) and 1% penicillin/streptomycin (100 U/ml; Fisher Scientific, 15140163). For co-culture differentiation experiments, OP9 cells were seeded into 96-well flat-bottomed plates (Genesee, 25-109). Once confluent, the cells were subjected to mitotic inactivation by irradiation at 300 Gy prior to co-culture with precursors.

### Generation of the isogenic iPSC line

CRISPR-Cas9 corrected line was generated by the Columbia Stem Cell Initiative Stem Cell core as previously described ^40^. Six validated synthetic guide RNAs (sgRNAs) were designed using the CRISPR Design Tool (https://design.synthego.com/#/, Synthego) to target two regions in the Homo Sapiens *GINS4* (NM_032336) exon7: chr8:41,541,835 region where the c.G511C (p.V171L) is located, and chr8:41,541,895 region where the c.C571T (p.Q191X) is located.

GTGGCGTGCATCTTTCACAGTGG

GATCTGGTTTGGGAACTGCCAGG

AGAACGACAAGAAAACATACTGG

GTGGCGTGCATCTTTCACAG

AGAACGACAAGAAAACATAC (used for editing)

CAAtGtGAttAGGAAAtCCt

All six sgRNAs were commercially synthesized (CRISPRevolution sgRNA EZ Kit, 1.5 nmol, Synthego) and used in combination with the Alt-R S.p. HiFi Cas9 Nuclease V3 (IDT, Integrated DNA Technologies) to assess their targeting efficiency by electroporation of the mutant GINS4 iPSC line using the 4D-Nucleofector (Lonza), P3 primary cell 4D-Nucleofector X kit S (Lonza) and program CA137. The Inference of CRISPR Edits (ICE) online tool (Synthego) was used to determine cleavage efficiency, then sgRNA 5 was chosen because of its cutting efficiency (44%) and better off-target score. The single-stranded oligodeoxynucleotide (ssODN) CAGATCTAGATTCTTAC***G***TGTTTCTGAGAGTGAGAGAACGACAAGAAAACATACTc

GTAGAACCAGACACAGATGAG***C***AGAGGTGAGTGGCGTGCATCTT (bold cases indicate target corrections; lower case indicates PAM mutation) was designed as a complementary sequence to these sgRNA target strands and purchased (IDT), then 15 μg ssODN, 15 μg sgRNA and 10 μg Cas9 were incubated for 25 minutes at RT to generate the RNP complex that was mixed with 1×10^6^ cells. ICE analysis was used to determine knock-in efficiency. 48-72 hours after electroporation, cells were seeded at low-density (single-cell, 5 × 10^3^ cells in a 10-cm dish) then colonies were manually picked in a 96-well plate for genotyping. Correction was confirmed with Sanger sequencing. Five clones were heterozygote while three clones were homozygote, and the edited lines were karyotyped by G-banding.

### NK cell differentiation

NK cell differentiation from iPSC was performed as described in Knorr et al.^23^. Briefly, 9000 cells per well were plated on 96 well polystyrene round bottomed microwell plates (Fisher Scientific, 262162) for EB formation in APEL-2 media (StemCell Technologies, 5275) containing 10μM ROCK inhibitor, 40ng/ml SCF (Peprotech, 300-07), 20ng/ml BMP (R&D, 314-BP), 20ng/ml VEGF (R&D, 293-VE), and 1μM GSK inhibitor (CHIR99021; StemCell Technologies, 72052). EBs were cultured for six days. For spin-EB differentiation, day 6 EBs were transferred to gelatin (Millipore, ES-006-B) coated 24 well plates (4EB/well; Fisher Scientific, 08-772-1C) with NK cell differentiation basal media supplemented with 5ng/ml IL-3 (1st week only; Peprotech, 200-03), 10ng/ml IL-15 (Peprotech, 200-15), 10ng/ml Flt3L (Peprotech, 300-19), 20ng/ml SCF, and 20ng/ml IL-7 (Peprotech, 200-07). NK cell differentiation basal media is made with 55% DMEM (Fisher Scientific, 11965118), 28% Hems F12 (Fisher Scientific, 10-080-CV), 15% human AB serum (GeminiBio, 100-512), 1% L-Glutamax (Fisher Scientific, 35050079), 1% Penicillin/streptomycin (100U/ml), 25μM BME (Fisher Scientific, 21985023), 50 μM Ethanolamine (Sigma, E0135-100ML), 25 μg/mL Ascorbic acid (Sigma, A5960-25G), and 5 ng/mL sodium selenite (Sigma, S5261-25G). After transfer, the cells from EB propagate and differentiate to NK cells in the 24 well plate for 35 days with weekly half media change.

For OP9 supported differentiation, day 6 EBs were dissociated with TrypLE (Fisher Scientific, 12605028), and CD34^+^ cells were sorted (BD FACS Aria II). 2 × 10^3^ CD34^+^ cells were seeded on each well of 96 well plates with irradiated OP9 (ATCC) stroma. For differentiation, the same differentiation basal media and cytokine supplements as differentiation supported by autologous stroma were used for 35 days with weekly half media change. T cell differentiation was performed using STEMDiff T cell Kit (StemCell Technologies, 100-0194) according to the manufacturer’s protocol. Sorted CD34^+^ HSC or CD117^+^ were differentiated on irradiated OP9 with the same differentiation basal media and cytokine supplements as differentiation supported by autologous stroma were used for 35 days with weekly half media change. Cells were treated with aphidicolin as stated in the text.

### CFU assay

Methods were adapted from Dege et al.^41^ for CFU assays. 9000 cells were seeded in each well of round bottom 96 well plates for EB formation in APEL-2 media with 10 ng/ml BMP4. On day1, bFGF (final concentration 5 ng/ml; RnD, 233-FB-025) was added. 18 hours later, final concentrations of either 3 μM CHIR99021 for definitive hematopoietic differentiation or 3 μM IWP2 (Selleckchem, S7085) and 1 ng/ml activin A (StemCell Technologies, 78001) for primitive hematopoietic differentiation were added to the media for 30 hours. EBs were placed in APEL-2 media with 15ng/ml VEGF and 5ng/ml bFGF for 3 days. On day 6, final concentrations of 100ng/ml SCF, 2IU/ml EPO (RnD, 287-TC-500), 10ng/mL IL-6 (Peprotech, 200-06), 5ng/ml IL-11 (Peprotech, 200-11), and 25ng/ml IGF-1 (Peprotech, 100-11) were added to the culture media. EBs were cultured in 37°C, 5% CO2, and 5% O2 condition. On day 8, CHIR99021 treated EBs were dissociated with TrypLE and CD34^+^CD43^−^CD73^−^CD184^−^ population was sorted and 10^4^ cells/well were plated on low 96-well low-adherent cell culture plates (Fisher Scientific, 07-200-603) in HE media (APEL-2 with 5ng/ml VEGF, 5ng/ml bFGF, 100ng/ml SCF, 2IU/ml EPO, 10ng/ml IL-6, 5ng/ml IL-11, 25ng/ml IGF-1, 30ng/ml TPO (Peprotech, 300-18), 10ng/ml BMP4, 20ng/ml SHH (RnD, 1845-SH-025), 10μg/ml angiotensin II (RnD, 1158/5), 100μM losartan potassium (RnD, 3798/50)). Once the cells aggregate into clumps, they were transferred to Matrigel coated 24 well plate in HE media. On day 20, CD34^+^CD45^+^ population was isolated with FACS sort.

On day 8, IWP2 treated EBs were put on day8 media (APEL-2 with 100ng/ml SCF, 2UI/ml EPO, 10ng/ml IL-6, 5ng/ml IL-11, 25ng/ml IGF-1, 30ng/ml TPO, 10ng/ml Flt-3L, and 30ng/ml IL-3) for two days. Then on day 10, CD43^+^CD34^−^ population was isolated with FACS. Sorted cells were plated in Methocult optimum media (2-3 × 10^3^ cells/well; StemCell Technologies, H4034) according to manufacturer’s protocol. Colonies were counted after 7 - 10 days.

### Flow cytometric analysis and sorting

Flow cytometry was performed at stated time points using antibodies listed in Supplementary Table 1. Cells were dissociated with TrypLE and then incubated with surface receptor antibodies and a fixable viability dye. After a wash, cells were fixed and permeabilized using CytoFix/CytoPerm (Fisher Scientific, BDB554714) or Foxp3/Transcription factor staining buffer (Thermo Fisher, 00-5523-00) according to manufacturer’s protocol then cells were incubated with intracellular antibodies. For EdU staining, cells were incubated with 10μM EdU for 2h before harvesting, and the signals were detected using Click-iT EdU Alexa Fluor 488 Flow Cytometry Assay Kit (Thermo Fisher, C10420) according to manufacturer’s protocol. Data were acquired on a Bio-Rad ZE5 Cell Analyzer and analyzed using FlowJo 10 (RRID: SCR_014583, BD Biosciences) or OMIQ (Dotmatics). Detailed gating strategies for experiments are represented in each figure. Student’s T-test was used to compare frequencies of populations between cell lines and p-values are indicated in each figure legend. For all iPSC experiments, cells were selected based on FSC and SSC. FSC-H and FSC-A were plotted against each other to select single cells, and then Zombie^neg^ cells were selected as live cells. For PBMC experiments, lymphocytes were selected based on FSC and SSC. FSC-H and FSC-A were plotted against each other to select single cells, and then propidium iodide (PI)^-^ cells were selected as live cells.

For sorting, dissociated cells or PBMC isolated from fresh cord blood with density centrifugation were incubated with surface receptor antibodies. After a wash, cells were resuspended in PBS with PI (1:200), and the desired populations were sorted on a BD Aria II cytometer with a 100 µm nozzle. Sorted cells were cultured directly after isolation.

### RNA-seq

Dissociated cells from iPSC cultures were stained with anti-CD34 (RRID:AB_1732005, Biolegend, 343604) or anti-CD45 (RRID:AB_314404, Biolegend, 304016) antibody and propidium iodide (PI). PI^−^CD34^+^, PI^−^CD34^−^, PI^−^CD45^+^, and PI^−^CD45^−^ cells were sorted by flow cytometry with BD Aria II cytometer. For isolation of primary samples, PBMCs were stained with PE-Cy7 CD34 (RRID:AB_2629726, Biolegend, 343616), APC CD14 (RRID:AB_314190, Biolegend, 301808), Pacific Blue CD19 (RRID:AB_2073118, Biolegend, 302232), Alexa488 CD56 (RRID:AB_2564093, Biolegend, 362518), BUV CD3 BD (RRID:AB_2870222, Biosciences, 612940). T cells, NK cells, CD34+ cells and B cells were isolated and directly processed. 2∼10 × 10^4^ cells were lysed, and RNA was collected with NucleoSpin RNA XS kit (Takara, 740902) according to the manufacturer’s protocol. RNA quality control was performed using the Agilent 2100 Bioanalyzer System. Libraries were generated with SMARTer stranded RNA-Seq kit (Takara, 634839) and sequenced on NovaSeq 6000 (Illumina). The FASTQ files were generated using FastQC (RRID: SCR_014583, v0.11.7) and trimmed reads (RRID: SCR_011848, Trimmomatic 0.38) were mapped to UCSC hg38 with HISAT2 (RRID:SCR_015530, v2.1.0, Bowtie2, RRID:SCR_016368, v2.3.4.1) splice-aware aligner. The sequencing depth is between 58 and 80 million mappable reads. Transcript is assembled by StringTie (RRID: SCR_016323, v2.1.3b) and abundance of genes are calculated as read counts and FPKM. CoGAPS (RRID: SCR_001479, v3.16.0, https://github.com/FertigLab/CoGAPS) was run on log2 RNA-seq FPKM data for a range of k patterns (k=15 selected) and uncertainty as 10%^28,42^. Stable solution for 15 patterns were calculated based on 4 sets (set in setDistributedParams(nSets=4)), and nIterations of 50,000. The weights of each gene in each pattern are saved and ordered in a p-matrix. Gene ontology terms were identified using PANTHER (RRID: SCR_004869, v. 18.0), homosapience, biological process options. CoGAPS patterns were projected to other datasets using the projectR function in the projectR package (v1.12.0, https://github.com/genesofeve/projectR/) ^43^. Briefly, both datasets were sub-setted using only the common genes between them. Projections were calculated by matrix multiplication between the weights of our 15 CoGAPS patterns and gene expression values of the public RNA-seq datasets. This calculation resulted in projected values for each sample or single cell for each pattern. Bone marrow atlas data was plotted with tSNE calculated on the summary data using Rtsne function from Rtsne package (RRID: SCR_061900, v.0.6) ^29^. Projected values were represented by color intensity. Sample identities, previously defined in the original publications, were used to name clusters in both datasets. All RNA-seq data and related public data are accessible using the gEAR/NeMO Analytics framework ^44,45^ at https://nemoanalytics.org/p?l=NKseoEtAl2023. All gene expression analysis datasets are available in the Gene Expression Omnibus (GEO, RRID:SCR_005012) under the accession number GSE267366 and GSE288864. To assess allelic bias, nucleotide-level counts at both of the GINS4 variants, c.G511 (p.171) and c.C571 (p.191), were obtained from BAM files using IGV (RRID:SCR_011793, v. 2.8.2). Pie charts were generated using count numbers.

### Western blot

Samples were collected and sonicated in RIPA buffer (Fisher Scientific, 89900) with protease inhibitor (Fisher Scientific, 78443). The protein concentration was measured with Pierce BCA protein assay kit (Fisher Scientific, 23225). The same amount of protein was loaded in each well. The samples were run on 12% Bis-Tris gels and transferred onto nitrocellulose membranes (Fisher Scientific, LC2000). The membranes were blocked and incubated with primary antibody overnight at 4°C. After secondary antibody incubation for 30 min at room temperature, the blots were imaged with Li-COR and analyzed with ImageJ (RRID: SCR_003070, v. 2.0.0-rc-69). The Western blot bands were quantified using the ImageJ (RRID: SCR_003070, v. 2.0.0-rc-69) ‘plot lanes’ function. All bands were normalized to actin or tubulin loading controls that are used on the same blot. Student T-test was used to compare the protein expression levels.

### Digital PCR

Alle-specific gene expression was measured using QuantStudio Absolute Q Digital PCR System (Thermo Fisher) according to the manufacturer’s instruction. Custom TaqMan SNP genotyping probes for two *GINS4* variant and control *NOC2L* were designed based on a target nucleotide (reference-VIC/alternative-FAM) with 44 base pairs of left and right arms. Due to limited access to patient blood, and low NK cell counts in some of the samples, we pooled 3-4 vials of PBMC collected at two different time points to generate a single sample per person and cell type. mRNA was isolated from sorted cells T, NK, and B cells using NucleoSpin Kit (Takara) followed by cDNA library synthesis (SMART-Seq mRNA LP, Takara). 10 µl of PCR mixture was prepared with 10 ng of cDNA, 2 µl of 5x PCR master mix, and 0.25 µl of 40x probe/primer mix. 9 µl of PCR mixture loaded into the plate and covered with 15 µl of isolation buffer and gaskets. QC was checked with ROX signal from more than 20,000 chambers. Pie charts were generated based on the concentration of each allele per 1ul of RNA.

Probes: Left arm-[VIC/FAM]-Right arm

*GINS4* V171L

TGGACCTCTTTCGGGCAGTTCCCAAACCAGATCTAGATTCTTAC[G/C]TGTTTCTGA GAGTGAGAGAACGACAAGAAAACATACTGGTAGAA

*GINS4* Q191X

GAGAACGACAAGAAAACATACTGGTAGAACCAGACACAGATGAG[C/T]AGAGGGAC TACGTGATTGACCTGGAGAAGGGCTCACAGCACTTG

*NOC2L* Hg38_ Chr1:946247

NM_015658.4:exon16:c. 1843 G>A:p.L615L

CAGTGGAAGCCTGGGAGAAGCTGACCCGGGAAGAGGGGACACCC[C/T]TGACCTT GTACTACAGCCACTGGCGCAAGCTGCGTGACCGGGAG

## Acknowledgments

We would like to thank the staff of the Columbia Stem Cell Initiative Flow Cytometry Core Facility, under the leadership of Michael Kissner, at Columbia University Irving Medical Center for their contributions to the work presented in this manuscript. We thank Dr. Dusan Bogunovic for helpful feedback and Dr. Jordan Orange for sharing GINS4 iPSC lines.

## Data and materials availability

All RNA-seq data and related public data are accessible using the gEAR/NeMO Analytics framework ^44,45^ at https://nemoanalytics.org/p?l=NKseoEtAl2023. All gene expression analysis datasets are available in the Gene Expression Omnibus (GEO) under the accession numbers GSE267366 and GSE288864.

## Supplementary Figure Legends

**Figure S1.**
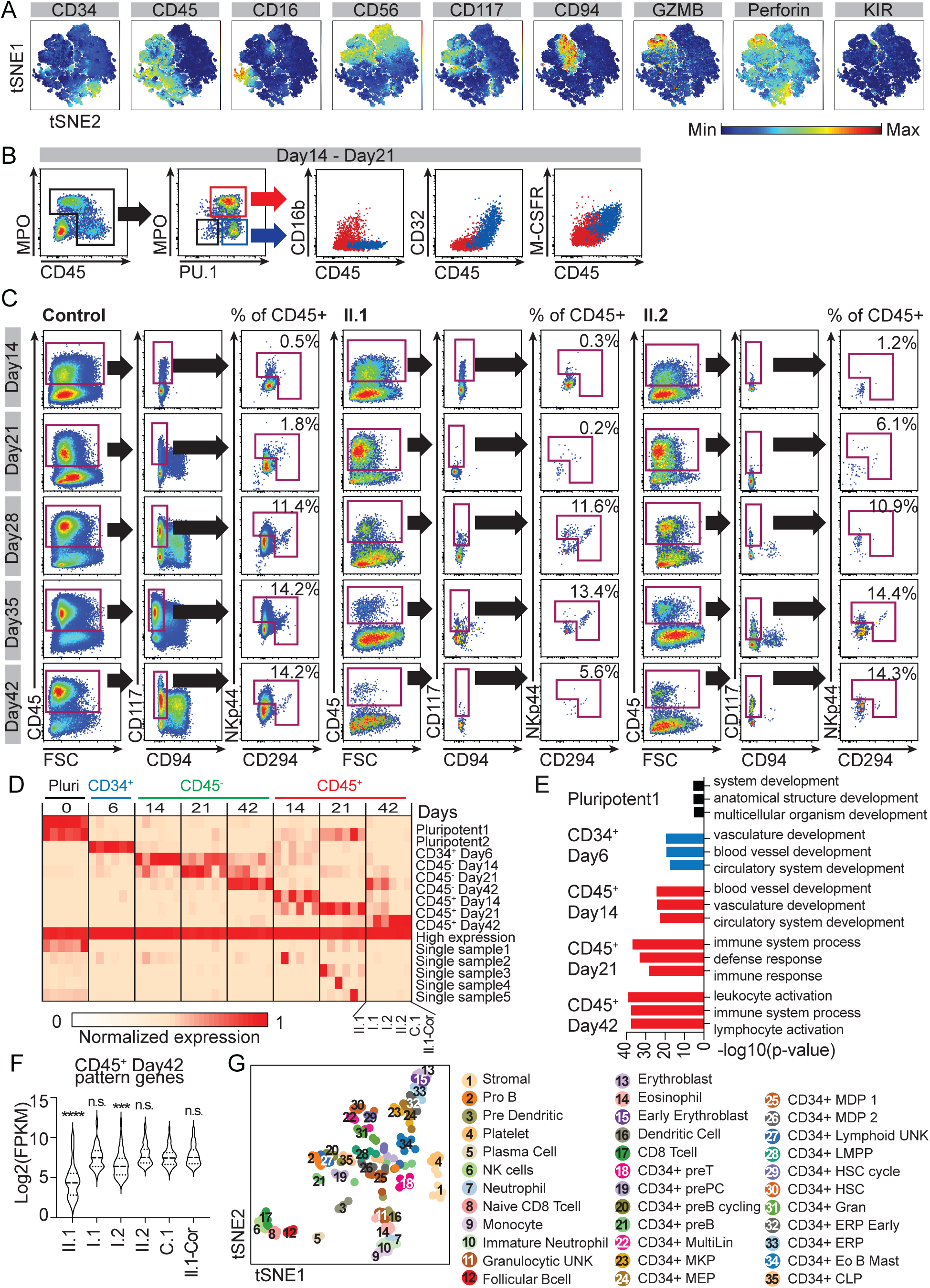
Autologous stroma differentiation using the spin-EB method and transcriptional analysis. NK cells were differentiated from iPSC lines. Flow cytometry was performed to monitor the progression of NK cell differentiation and identities of non-NK CD45^+^ cells. Bulk RNA-Seq was performed on differentiating iPSCs to assess transcriptional changes. **(A)** tSNE plot summarizing flow cytometry data from six lines (I.1, I.2, II.1, II.2, C.1, and II.1-Cor) colored by each marker. **(B)** Representative flow cytometry results showing additional markers for the myeloid identity of non-NK cell CD45^+^ populations on days 14 and 21. The gating strategy shows selection of CD45^dim^ and CD45^+^ populations. MPO^+^CD45^dim^PU.1^+^ cells in red box are represented as red dots, and MPO^−^CD45^+^PU.1^+^ cells in blue box are represented as blue dots in CD16b, CD32, and M-CSFR (CD115) plots. **(C)** Representative flow cytometry plots demonstrating emergence of ILC-like cells on days 14 and 42 in control, II.1 and II.2 lines. The gating strategy shows selection of CD45^+^CD117^+^CD94^−^ populations at each time points. Freqeuncy of NKp44^+^ or CD294^+^ ILC-like cells are calculated as % of CD45^+^ cells. **(D)** Heatmap representing the 15 CoGAPS patterns detected from all samples. Each pattern is named based on the condition with the highest expression of the signature. **(E)** GO terms in hematopoietic lineage related patterns (3 picked from the top 6 GO terms with the lowest p-values). **(F)** Expression levels of top 200 genes in NK related CD45^+^ Day42 pattern showing significantly low expression of this pattern in II.1 and I.2 compared to the C.1 line (student t test, ***p<0.005, ****p<0.001). **(G)** tSNE generated with complete annotation from the original manuscript ^29^.

**Figure S2.**
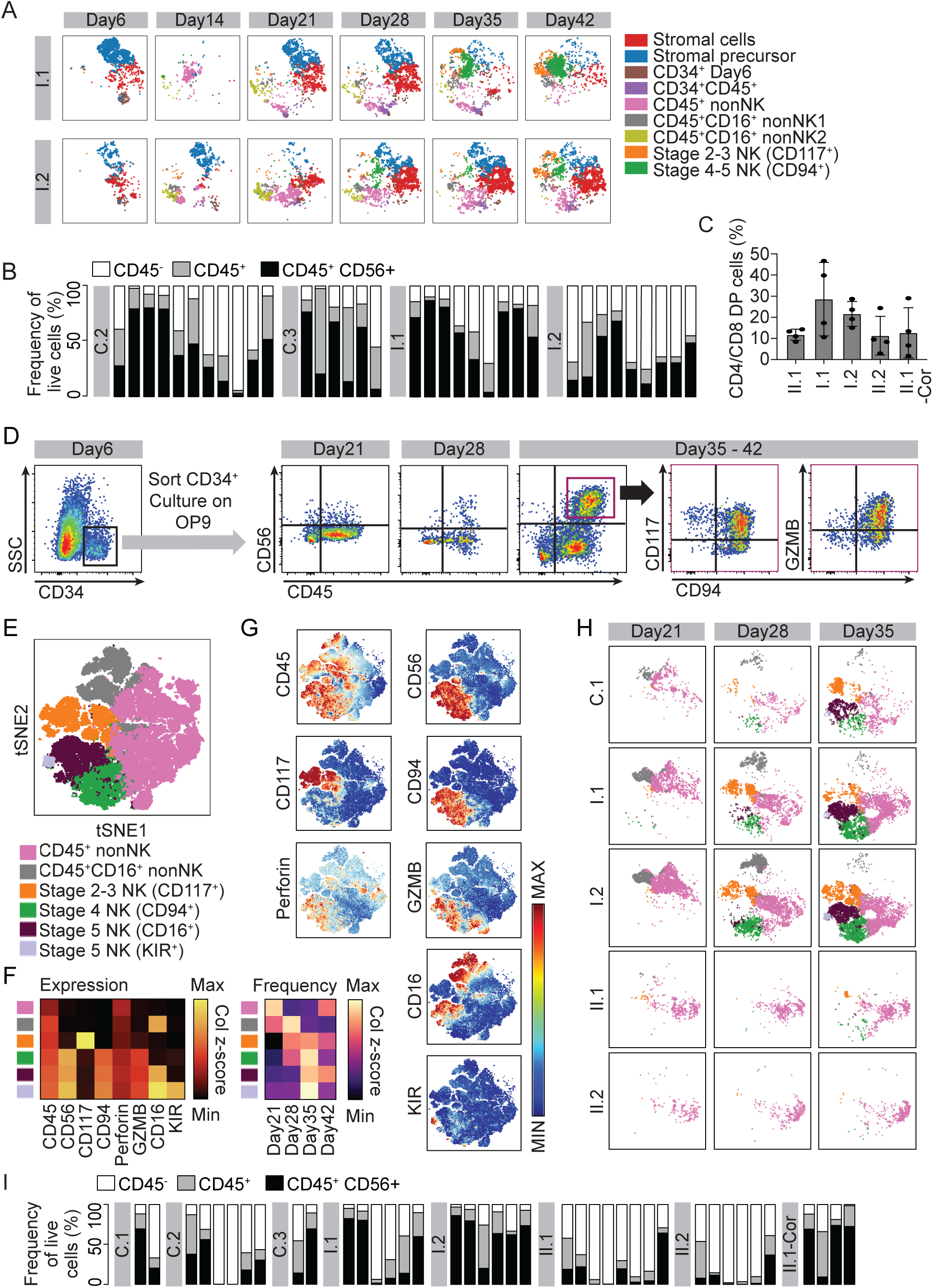
Different lines have distinct NK cell potentials. Hematopoietic potentials were tested on multiple iPSC lines with autologous stromal cells or by sorting CD34^+^ cells at day 6 and co-culturing with OP9 stroma. Flow cytometry was performed to monitor the progression of differentiation, and the data were concatenated, normalized, and visualized. **(A)** Parsed tSNE showing changes in cell populations in different lines from days 6 to 42 from lines I.1 and I.2 on autologous stroma. **(B)** Stacked bar graphs show the population of cells with hematopoietic (CD45) and NK cell (CD56) markers on days 35 or 42 of each autologous stroma experiment from C.2, C.3, I.1, and I.2 lines (n=6-11). **(C)** Frequency of CD4^+^ CD8^+^ double-positive T cells arising from each cell line (n=4 technical replicates, error bars: SD). **(D)** Dot plots showing a representative differentiation of the control C.1 line. On day 6, CD34^+^ cells were sorted and plated on OP9. CD45, CD56, CD94, and GZMB positive populations emerge between days 21 and 42. **(E)** tSNE plot of summarizing flow cytometry data of a representative OP9 experiment (n=2-8 technical replicates) from five lines (C.1, I.1, I.2, II.1, and II.2) at 4 weekly time points beginning on day 21 (14 days after CD34^+^ cell seeding). Each dot represents a single cell, and the cells are clustered into six groups. **(F)** Heatmaps showing a summary of relative protein expression patterns for each cluster (left) and temporal changes of each cluster (% of cells in each cluster from all live cells, right). **(G)** tSNE plots colored by each protein expression level represent the cluster identities. **(H)** Parsed tSNE plots showing cell populations on days 21, 28, and 35 for each cell line revealing NK cell deficiencies recapitulated in the GINS4 variant lines on OP9 (II.1 and II.2). **(I)** Stacked bar graphs show the population of cells with hematopoietic (CD45) and NK cell (CD56) markers on days 35 or 42 of each OP9 experiment from C.1, C.2, C.3, I.1, I.2, II.1, II.2, and II.1-Cor lines (n=2-8).

**Figure S3.**
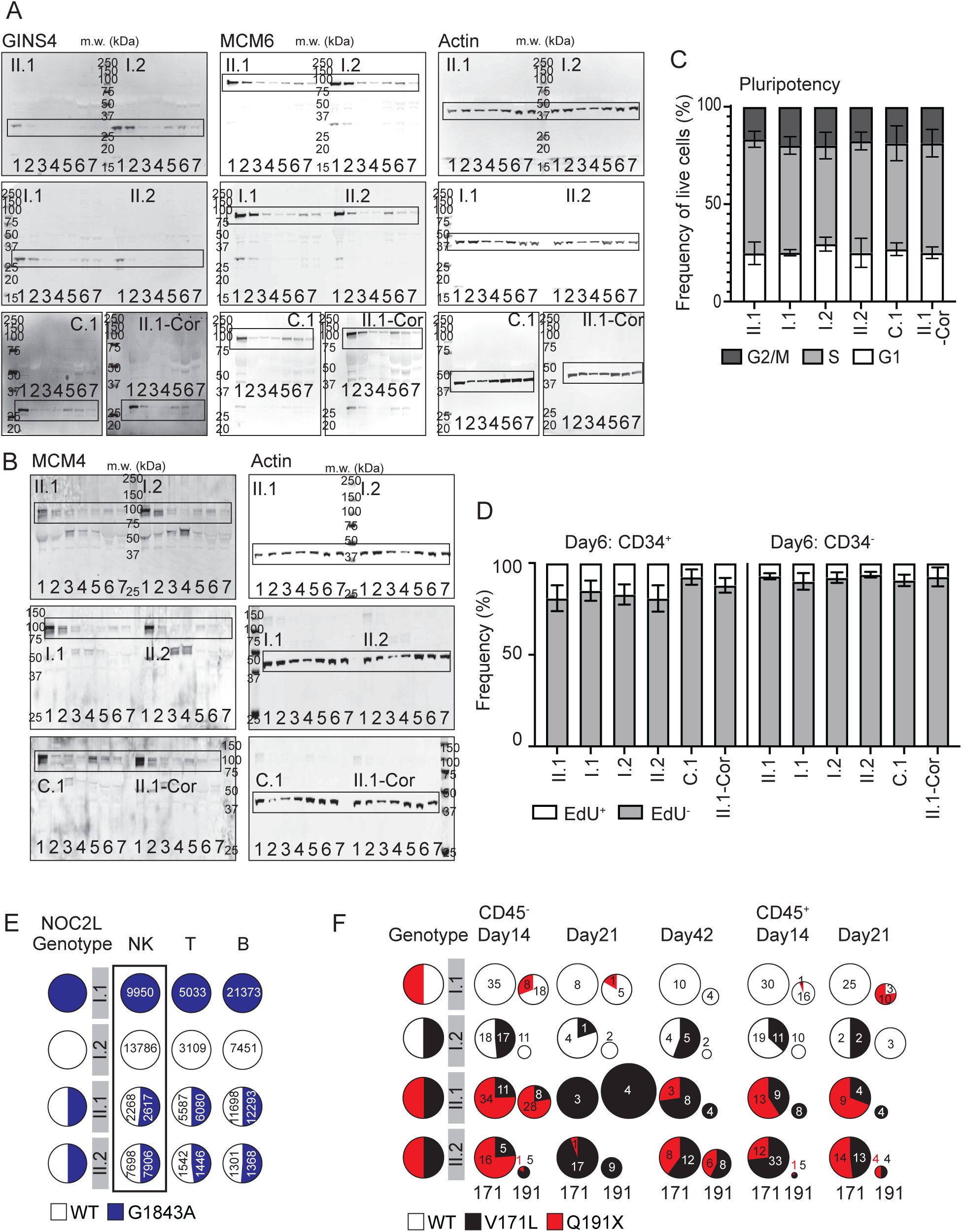
Full Western blots and additional flow cytometry analyses show cell cycle progression. Western blots were performed to understand the dynamics of cell cycle related proteins. Barplots and tSNE show additional flow cytometry analyses for cell cycle and NK differentiation efficiency. **(A)** Full western blots of GINS4, MCM6, and Actin in six lines (I.1, I.2, II.1, II.2, C.1, and II.1-Cor) at all time points from autologous stroma experiments. Lanes 1-7 represent 0, 6, 14, 21, 28, 35, and 42 days, respectively. **(B)** Full western blots of MCM4, and Actin in six lines (I.1, I.2, II.1, II.2, C.1, and II.1-Cor) at all time points. Lanes 1-7 represent 0, 6, 14, 21, 28, 35, and 42 days from autologous stroma experiments, respectively. **(C)** Cell cycle profiles of six iPSC lines measured by DAPI and EdU staining by flow cytometry (n=3-5 technical replicates, error bars=SD). **(D)** Day 6 cells from the EBs were stained with CD34 and EdU for assessing frequency of cells in the S phase by flow cytometry. Frequency of EdU^+^ populations in CD34^+^ or CD34^−^ populations from six cell lines (n=4 technical replicates, error bars=SD). **(E)** Digital PCR result of *NOC2L* gene in I.1, I.2, II.1, and II.2 lines represented in the pie charts show biallelic expression of the gene within sorted lymphocytes. Tested SNP is NM_015658.4:exon16:c. 1843 G>A:p.L615L. Numbers in the pie charts show concentration (copy/ul) per allele. **(F)** Pie charts show the contribution of each *GINS4* allele in I.1, I.2, II.1, and II.2 lines at different timepoints and subsets in iPSC differentiation samples on autologous stroma. Size of the pie charts represent relative counts at the p.191 locus compared to p.171 locus in each sample. Numbers in the pie charts show counts per allele.

## References

1. Tangye, S.G., Al-Herz, W., Bousfiha, A., Cunningham-Rundles, C., Franco, J.L., Holland, S.M., Klein, C., Morio, T., Oksenhendler, E., Picard, C., et al. (2022). Human Inborn Errors of Immunity: 2022 Update on the Classification from the International Union of Immunological Societies Expert Committee. J Clin Immunol 42, 1473–1507. 10.1007/s10875-022-01289-3.

2. Gruber, C., and Bogunovic, D. (2020). Incomplete penetrance in primary immunodeficiency: a skeleton in the closet. Hum Genet 139, 745–757. 10.1007/s00439-020-02131-9.

3. Miyazawa, H., and Wada, T. (2021). Reversion Mosaicism in Primary Immunodeficiency Diseases. Front Immunol 12, 783022. 10.3389/fimmu.2021.783022.

4. Staels, F., Collignon, T., Betrains, A., Gerbaux, M., Willemsen, M., Humblet-Baron, S., Liston, A., Vanderschueren, S., and Schrijvers, R. (2021). Monogenic Adult-Onset Inborn Errors of Immunity. Front Immunol 12, 753978. 10.3389/fimmu.2021.753978.

5. Kaneko, H., Kawamoto, N., Asano, T., Mabuchi, Y., Horikoshi, H., Teramoto, T., Matsui, E., Kondo, M., Fukao, T., Kasahara, K., and Kondo, N. (2005). Leaky phenotype of X-linked agammaglobulinaemia in a Japanese family. Clin Exp Immunol 140, 520–523. 10.1111/j.1365-2249.2005.02784.x.

6. Schwab, C., Gabrysch, A., Olbrich, P., Patino, V., Warnatz, K., Wolff, D., Hoshino, A., Kobayashi, M., Imai, K., Takagi, M., et al. (2018). Phenotype, penetrance, and treatment of 133 cytotoxic T-lymphocyte antigen 4-insufficient subjects. J Allergy Clin Immunol 142, 1932–1946. 10.1016/j.jaci.2018.02.055.

7. Gineau, L., Cognet, C., Kara, N., Lach, F.P., Dunne, J., Veturi, U., Picard, C., Trouillet, C., Eidenschenk, C., Aoufouchi, S., et al. (2012). Partial MCM4 deficiency in patients with growth retardation, adrenal insufficiency, and natural killer cell deficiency. J Clin Invest 122, 821–832. 10.1172/JCI61014.

8. Mace, E.M., Paust, S., Conte, M.I., Baxley, R.M., Schmit, M.M., Patil, S.L., Guilz, N.C., Mukherjee, M., Pezzi, A.E., Chmielowiec, J., et al. (2020). Human NK cell deficiency as a result of biallelic mutations in MCM10. J Clin Invest 130, 5272–5286. 10.1172/JCI134966.

9. Cottineau, J., Kottemann, M.C., Lach, F.P., Kang, Y.H., Vely, F., Deenick, E.K., Lazarov, T., Gineau, L., Wang, Y., Farina, A., et al. (2017). Inherited GINS1 deficiency underlies growth retardation along with neutropenia and NK cell deficiency. J Clin Invest 127, 1991–2006. 10.1172/JCI90727.

10. Conte, M.I., Poli, M.C., Taglialatela, A., Leuzzi, G., Chinn, I.K., Salinas, S.A., Rey-Jurado, E., Olivares, N., Veramendi-Espinoza, L., Ciccia, A., et al. (2022). Partial loss-of-function mutations in GINS4 lead to NK cell deficiency with neutropenia. JCI Insight 7. 10.1172/jci.insight.154948.

11. Hughes, C.R., Guasti, L., Meimaridou, E., Chuang, C.H., Schimenti, J.C., King, P.J., Costigan, C., Clark, A.J., and Metherell, L.A. (2012). MCM4 mutation causes adrenal failure, short stature, and natural killer cell deficiency in humans. J Clin Invest 122, 814–820. 10.1172/JCI60224.

12. Bernard, F., Picard, C., Cormier-Daire, V., Eidenschenk, C., Pinto, G., Bustamante, J.C., Jouanguy, E., Teillac-Hamel, D., Colomb, V., Funck-Brentano, I., et al. (2004). A novel developmental and immunodeficiency syndrome associated with intrauterine growth retardation and a lack of natural killer cells. Pediatrics 113, 136–141. 10.1542/peds.113.1.136.

13. Eidenschenk, C., Jouanguy, E., Alcais, A., Mention, J.J., Pasquier, B., Fleckenstein, I.M., Puel, A., Gineau, L., Carel, J.C., Vivier, E., et al. (2006). Familial NK cell deficiency associated with impaired IL-2- and IL-15-dependent survival of lymphocytes. J Immunol 177, 8835–8843. 10.4049/jimmunol.177.12.8835.

14. Eidenschenk, C., Dunne, J., Jouanguy, E., Fourlinnie, C., Gineau, L., Bacq, D., McMahon, C., Smith, O., Casanova, J.L., Abel, L., and Feighery, C. (2006). A novel primary immunodeficiency with specific natural-killer cell deficiency maps to the centromeric region of chromosome 8. Am J Hum Genet 78, 721–727. 10.1086/503269.

15. Fenwick, A.L., Kliszczak, M., Cooper, F., Murray, J., Sanchez-Pulido, L., Twigg, S.R., Goriely, A., McGowan, S.J., Miller, K.A., Taylor, I.B., et al. (2016). Mutations in CDC45, Encoding an Essential Component of the Pre-initiation Complex, Cause Meier-Gorlin Syndrome and Craniosynostosis. Am J Hum Genet 99, 125–138. 10.1016/j.ajhg.2016.05.019.

16. Guilz, N.C., Conte, M.I., Salinas, S.A., Chinn, I.K., Lupski, J.R., and Mace, E.M. (2023). CDC45 haploinsufficiency as a novel cause of natural killer cell deficiency. Clinical Immunology 250 (2023), 109356.

17. Lim, H.J., Jeon, Y., Jeon, C.H., Kim, J.H., and Lee, H. (2011). Targeted disruption of Mcm10 causes defective embryonic cell proliferation and early embryo lethality. Biochim Biophys Acta 1813, 1777–1783. 10.1016/j.bbamcr.2011.05.012.

18. Shima, N., Alcaraz, A., Liachko, I., Buske, T.R., Andrews, C.A., Munroe, R.J., Hartford, S.A., Tye, B.K., and Schimenti, J.C. (2007). A viable allele of Mcm4 causes chromosome instability and mammary adenocarcinomas in mice. Nat Genet 39, 93–98. 10.1038/ng1936.

19. Ueno, M., Itoh, M., Kong, L., Sugihara, K., Asano, M., and Takakura, N. (2005). PSF1 is essential for early embryogenesis in mice. Mol Cell Biol 25, 10528–10532. 10.1128/MCB.25.23.10528-10532.2005.

20. Mohri, T., Ueno, M., Nagahama, Y., Gong, Z.Y., Asano, M., Oshima, H., Oshima, M., Fujio, Y., and Takakura, N. (2013). Requirement of SLD5 for early embryogenesis. PLoS One 8, e78961. 10.1371/journal.pone.0078961.

21. Guilz, N.C., Ahn, Y.O., Seo, S., and Mace, E.M. (2023). Unwinding the Role of the CMG Helicase in Inborn Errors of Immunity. J Clin Immunol 43, 847–861. 10.1007/s10875-023-01437-3.

22. Guilz, N.C., Ahn, Y.O., Fatima, H., Pedroza, L.A., Seo, S., Soni, R.K., Wang, N., Egli, D., and Mace, E.M. (2024). Replication Stress in Activated Human NK Cells Induces Sensitivity to Apoptosis. J Immunol 213, 40–51. 10.4049/jimmunol.2300843.

23. Knorr, D.A., Ni, Z., Hermanson, D., Hexum, M.K., Bendzick, L., Cooper, L.J., Lee, D.A., and Kaufman, D.S. (2013). Clinical-scale derivation of natural killer cells from human pluripotent stem cells for cancer therapy. Stem Cells Transl Med 2, 274–283. 10.5966/sctm.2012-0084.

24. Scoville, S.D., Freud, A.G., and Caligiuri, M.A. (2017). Modeling Human Natural Killer Cell Development in the Era of Innate Lymphoid Cells. Front Immunol 8, 360. 10.3389/fimmu.2017.00360.

25. Hazenberg, M.D., and Spits, H. (2014). Human innate lymphoid cells. Blood 124, 700–709. 10.1182/blood-2013-11-427781.

26. Parodi, M., Favoreel, H., Candiano, G., Gaggero, S., Sivori, S., Mingari, M.C., Moretta, L., Vitale, M., and Cantoni, C. (2019). NKp44-NKp44 Ligand Interactions in the Regulation of Natural Killer Cells and Other Innate Lymphoid Cells in Humans. Front Immunol 10, 719. 10.3389/fimmu.2019.00719.

27. Verma, R., Er, J.Z., Pu, R.W., Sheik Mohamed, J., Soo, R.A., Muthiah, H.M., Tam, J.K.C., and Ding, J.L. (2020). Eomes Expression Defines Group 1 Innate Lymphoid Cells During Metastasis in Human and Mouse. Front Immunol 11, 1190. 10.3389/fimmu.2020.01190.

28. Fertig, E.J., Ding, J., Favorov, A.V., Parmigiani, G., and Ochs, M.F. (2010). CoGAPS: an R/C++ package to identify patterns and biological process activity in transcriptomic data. Bioinformatics 26, 2792–2793. 10.1093/bioinformatics/btq503.

29. Hay, S.B., Ferchen, K., Chetal, K., Grimes, H.L., and Salomonis, N. (2018). The Human Cell Atlas bone marrow single-cell interactive web portal. Exp Hematol 68, 51–61. 10.1016/j.exphem.2018.09.004.

30. Kilpinen, H., Goncalves, A., Leha, A., Afzal, V., Alasoo, K., Ashford, S., Bala, S., Bensaddek, D., Casale, F.P., Culley, O.J., et al. (2017). Common genetic variation drives molecular heterogeneity in human iPSCs. Nature 546, 370–375. 10.1038/nature22403.

31. Kim, S.K., Seo, S., Stein-O’Brien, G., Jaishankar, A., Ogawa, K., Micali, N., Luria, V., Karger, A., Wang, Y., Kim, H., et al. (2024). Individual variation in the emergence of anterior-to-posterior neural fates from human pluripotent stem cells. Stem Cell Reports 19, 1336–1350. 10.1016/j.stemcr.2024.07.004.

32. Herrera, L., Salcedo, J.M., Santos, S., Vesga, M.A., Borrego, F., and Eguizabal, C. (2017). OP9 Feeder Cells Are Superior to M2-10B4 Cells for the Generation of Mature and Functional Natural Killer Cells from Umbilical Cord Hematopoietic Progenitors. Front Immunol 8, 755. 10.3389/fimmu.2017.00755.

33. Stewart, O., Gruber, C., Randolph, H.E., Patel, R., Ramba, M., Calzoni, E., Huang, L.H., Levy, J., Buta, S., Lee, A., et al. (2025). Monoallelic expression can govern penetrance of inborn errors of immunity. Nature. 10.1038/s41586-024-08346-4.

34. Orr, S.J., Gaymes, T., Ladon, D., Chronis, C., Czepulkowski, B., Wang, R., Mufti, G.J., Marcotte, E.M., and Thomas, N.S. (2010). Reducing MCM levels in human primary T cells during the G(0)-->G(1) transition causes genomic instability during the first cell cycle. Oncogene 29, 3803–3814. 10.1038/onc.2010.138.

35. Xu, X., Hua, X., Brown, K., Ren, X., and Zhang, Z. (2022). Mcm2 promotes stem cell differentiation via its ability to bind H3-H4. Elife 11. 10.7554/eLife.80917.

36. Petryk, N., Dalby, M., Wenger, A., Stromme, C.B., Strandsby, A., Andersson, R., and Groth, A. (2018). MCM2 promotes symmetric inheritance of modified histones during DNA replication. Science 361, 1389–1392. 10.1126/science.aau0294.

37. Memar, N., Sherrard, R., Sethi, A., Fernandez, C.L., Schmidt, H., Lambie, E.J., Poole, R.J., Schnabel, R., and Conradt, B. (2024). The replicative helicase CMG is required for the divergence of cell fates during asymmetric cell division in vivo. Nat Commun 15, 9399. 10.1038/s41467-024-53715-2.

38. Carcamo-Orive, I., Hoffman, G.E., Cundiff, P., Beckmann, N.D., D’Souza, S.L., Knowles, J.W., Patel, A., Papatsenko, D., Abbasi, F., Reaven, G.M., et al. (2017). Analysis of Transcriptional Variability in a Large Human iPSC Library Reveals Genetic and Non-genetic Determinants of Heterogeneity. Cell Stem Cell 20, 518–532 e519. 10.1016/j.stem.2016.11.005.

39. Choi, J., Lee, S., Mallard, W., Clement, K., Tagliazucchi, G.M., Lim, H., Choi, I.Y., Ferrari, F., Tsankov, A.M., Pop, R., et al. (2015). A comparison of genetically matched cell lines reveals the equivalence of human iPSCs and ESCs. Nat Biotechnol 33, 1173–1181. 10.1038/nbt.3388.

40. Patel, A., Iannello, G., Diaz, A.G., Sirabella, D., Thaker, V., and Corneo, B. (2022). Efficient Cas9-based Genome Editing Using CRISPR Analysis Webtools in Severe Early-onset-obesity Patient-derived iPSCs. Curr Protoc 2, e519. 10.1002/cpz1.519.

41. Dege, C., and Sturgeon, C.M. (2017). Directed Differentiation of Primitive and Definitive Hematopoietic Progenitors from Human Pluripotent Stem Cells. J Vis Exp. 10.3791/55196.

42. Stein-O’Brien, G.L., Carey, J.L., Lee, W.S., Considine, M., Favorov, A.V., Flam, E., Guo, T., Li, S., Marchionni, L., Sherman, T., et al. (2017). PatternMarkers & GWCoGAPS for novel data-driven biomarkers via whole transcriptome NMF. Bioinformatics 33, 1892–1894. 10.1093/bioinformatics/btx058.

43. Sharma, G., Colantuoni, C., Goff, L.A., Fertig, E.J., and Stein-O’Brien, G. (2020). projectR: an R/Bioconductor package for transfer learning via PCA, NMF, correlation and clustering. Bioinformatics 36, 3592–3593. 10.1093/bioinformatics/btaa183.

44. Ament, S.A., Adkins, R.S., Carter, R., Chrysostomou, E., Colantuoni, C., Crabtree, J., Creasy, H.H., Degatano, K., Felix, V., Gandt, P., et al. (2023). The Neuroscience Multi-Omic Archive: a BRAIN Initiative resource for single-cell transcriptomic and epigenomic data from the mammalian brain. Nucleic Acids Res 51, D1075–D1085. 10.1093/nar/gkac962.

45. Orvis, J., Gottfried, B., Kancherla, J., Adkins, R.S., Song, Y., Dror, A.A., Olley, D., Rose, K., Chrysostomou, E., Kelly, M.C., et al. (2021). gEAR: Gene Expression Analysis Resource portal for community-driven, multi-omic data exploration. Nat Methods 18, 843–844. 10.1038/s41592-021-01200-9.

